# Network-based multi-omics integration reveals metabolic at-risk profile within treated HIV-infection

**DOI:** 10.1101/2022.06.08.495246

**Authors:** Flora Mikaeloff, Marco Gelpi, Rui Benfeitas, Andreas D. Knudsen, Beate Vestad, Julie Høgh, Johannes R. Hov, Thomas Benfield, Daniel Murray, Christian G Giske, Adil Mardinoglu, Marius Trøseid, Susanne D. Nielsen, Ujjwal Neogi

**Author notes:** Equal contribution (Susanne D. Nielsen, Ujjwal Neogi). **AUTHOR CONTRIBUTIONS** Conceptualization: UN, FM, RB; Clinical Study Design: SDN, TB, MG, ADK; Methodology: FM, MG, RB, ADK, AM, MT; Investigation, clinical: SDN, TB, MG, ADK; Investigation, computational and laboratory: FM, MG, RB, ADK, BV, JH, JRH; Visualization: FM, UN; Funding acquisition: UN, SDN; Project administration: SDN, RB, UN; Supervision: SDN, RB, UN; Writing – original draft: FM, UN; Writing – review & editing: MG, RB, ADK, BV, JH, JRH, TB, DM, CGG, MT, SDN. **Competing Interest Statement:** The authors declare no competing interests.

## Abstract

Multiomics technologies improve the biological understanding of health status in people living with HIV on antiretroviral therapy (PLWH_ART_). Still, a systematic and in-depth characterization of metabolic risk profile during successful long-term treatment is lacking. Here, we used multi-omics (plasma lipidomic and metabolomic, and fecal 16s microbiome) data-driven stratification and characterization to identify the metabolic at-risk profile within PLWH_ART_. Through network analysis and similarity network fusion (SNF), we identified three groups of PLWH_ART_ (SNF-1 to 3). The PLWH_ART_ at SNF-2 (45%) was a severe at-risk metabolic profile with increased visceral adipose tissue, BMI, higher incidence of metabolic syndrome (MetS), and increased di- and triglycerides despite having higher CD4^+^ T-cell counts than the other two clusters. However, the healthy-like and severe at-risk group had a similar metabolic profile differing from HC, with dysregulation of amino acid metabolism. At the microbiome profile, the healthy-like group had a lower α-diversity, a lower proportion of MSM, and was enriched in Bacteroides. In contrast, in at-risk groups, there was an increase in *Prevotella*, with a high proportion of men who have sex with men (MSM) confirming the influence of sexual orientation on the microbiome profile The multi-omics integrative analysis reveals a complex microbial interplay by microbiome-derived metabolites in PLWH_ART_. PLWH_ART_ those are severely at-risk clusters may benefit from personalized medicine and lifestyle intervention to improve their metabolic profile.

**Significance:** The network and factorization-based integrative analysis of plasma metabolomics, lipidomics, and microbiome profile identified three different diseases’ state -omics phenotypes within PLWH_ART_ driven by metabolomics, lipidomics, and microbiome that a single omics or clinical feature could not explain. The severe at-risk group has a dysregulated metabolic profile that potentiates metabolic diseases that could be barriers to healthy aging. The at-risk group may benefit from personalized medicine and lifestyle intervention to improve their metabolic profile.

## Introduction

Antiretroviral therapy (ART) has improved the immune profile by suppressing viral replication and reducing the morbidity and mortality of people living with HIV (PLWH). Yet living with HIV under ART induces a strong metabolic perturbation in the body due to virus persistence, immune activation, chronic low-grade inflammation, and treatment toxicity, mostly with older antiretrovirals ^1^. The biological shifts due to a mixed effect of drugs and viruses are also highly personalized depending on the patient genetic background, age, gender, immunological, and lifestyle factors ^2^. The long-term HIV infection, even with successful ART, is associated with an accentuated onset of non-AIDS-related comorbidities ^3^. Consequently, diseases of the aged population appear in relatively young HIV patients, including cardiovascular disease, liver-kidney disease, and neurocognitive and metabolic disorders ^4^.

Systems biological analyses are valuable methodologies for systematically understanding pathology and identifying potential novel treatment strategies ^5^. Microbiome studies provided enormous knowledge about the microbial association with the HIV status, sexual practice, and gender ^6-8^ and the possible interplay between HIV-related gut microbiota, immune dysfunction, and comorbidities like metabolic syndrome (MetS) and visceral adipose tissue (VAT) accumulation ^7^. Our extensive metabolomics studies from three different cohorts from India ^9^, Cameroon ^10^, and Denmark ^11^ with more than 500 PLWH indicated that disrupted amino acid (AA) metabolism in PLWH with ART (PLWH_ART_) following prolonged ART that plays the central role in the comorbidities such as MetS ^11^.

Multi-omic characterizations may offer insights into understanding the mechanisms underlying biological processes in a specific disease condition. The application of integrative omics to understand the disease pathogenesis in PLWH under suppressive ART is lacking. To the best of our knowledge, no integrative omics studies have been performed to understand complex biological phenotypes in PLWH during prolonged suppressive ART (PLWH_ART_). A recent longitudinal study integrating metabolomics, plasma protein biomarkers, and transcriptomics in patients’ samples identified potential lipid and amino acid metabolism perturbations in PLWH with immune reconstitution inflammatory syndrome (IRIS) ^12^. Our recent network-based integrative plasma lipidomics, metabolic biomarker, and clinical data indicated a coordinated role of clinical parameters like accumulation of visceral adipose tissue (VAT) and exposure to earlier generations of antiretrovirals with glycerolipids and glutamate metabolism in the pathogenesis of PLWH with MetS ^13^.

The present study aimed to identify the molecular data-driven phenotypic patient stratification using network-based integration of plasma metabolomics/lipidomics and fecal microbiota in a cohort of PLWH_ART_ with prolonged suppressive therapy to identify the at-risk metabolic profile following long-term successful therapy. We further investigated the underlying factors differing from these profiles and the link to their clinical phenotype to clarify risk factors for metabolic disease further.

## Results

### Comprehensive multi-omics characterization of PLWH on successful Cart

In this study, we used untargeted plasma metabolomics (877 metabolites) ^11^, lipidomics (977 lipids) ^13^, and fecal 16s rRNA microbiome [241 operational taxonomic unit (OTU)] data ^7^ from 97 PLWH_ART_ from the Copenhagen Comorbidity (COCOMO) cohort ^14^ where we have three levels of omics data available. Additionally, we included 48 clinical and demographical features comprising lifestyle habits (food, medicine, alcohol, smoking), comorbidities linked to obesity, and HIV-related measurements (viral load, treatment history, CD4 T-cell count, CD8 T-cell counts) (Table S1). The PLWH were mainly male (86%; 84/97), of Caucasian ethnic origin (81%, 79/97) with a median (IQR) age of 54 (48-63) years. The median (IQR) duration of the treatment was 15 (9-18) years. At the time of sample collection, the viral load was below detection level with successful immune reconstitution [median (IQR) CD4 T-cell count 713 (570-900) cells/μL]. Additionally, 20 HIV-negative controls (HC) with similar sex proportions (90%, 18/20) and median age (IQR) of 56 (50-67) years were used to reference multi-omics.

### Integrative omics-based similarity network fusion (SNF) identifies three clusters in PLWH_ART_

To stratify the PLWH_ART_ based on their molecular signature, we used Similarity Network Fusion (SNF) that constructs similarity matrices and networks of PLWH_ART_ for each of the omics and fuses into one network that represents the full spectrum of the underlying data and disease status in PLWH_ART_ ^15^. We identified three clusters of patients, defined as SNF-1 (N=19), SNF-2 (N=44), and SNF-3 (N=34) (Fig 1A). The concordance matrix based on Normalized Mutual Information (NMI) score (0=no mutual information, 1=perfect correlation) showed that lipids had the most influence in the final network (NMI=0.6), followed by metabolites (NMI=0.4) and finally, microbiome (NMI=0.3) (Fig 1B). Clear segregation of the SNF clusters (Fig 1C) was observed in the PCA based on the fused network values (Fig 1D) and PCA of single omics for lipidomics and metabolomics but not microbiome (Fig S1). The addition of HC with the clusters showed that the SNF-3 had an HC-like profile in metabolomics and lipidomics PCA plots (Fig S1).

**Figure 1:**
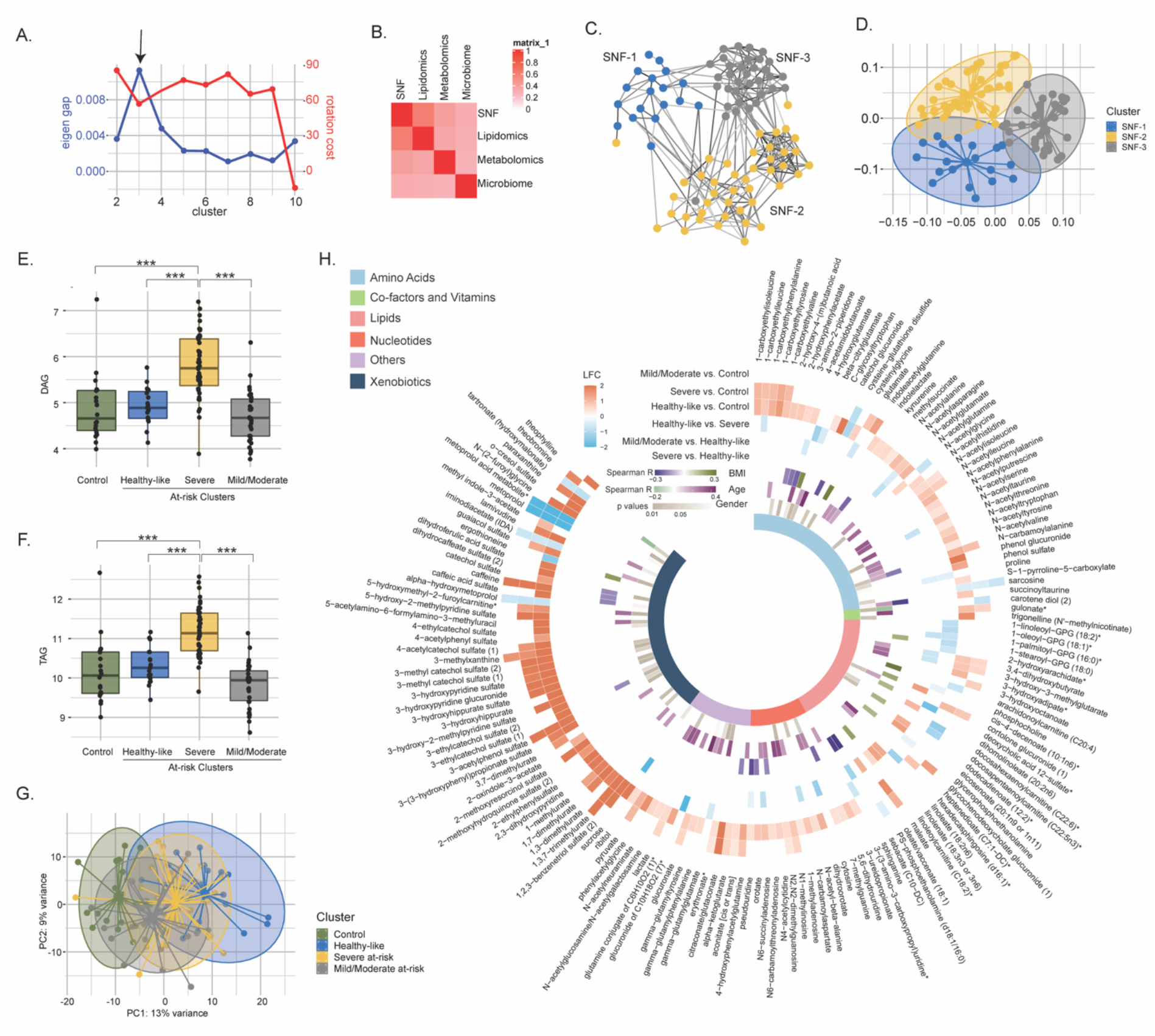
Similarity network fusion based on lipidomics, metabolomics, and microbiome integration. **(a)** Scatter plot showing the maximization of Eigen gap and the minimization of rotation cost for the optimization of the number of clusters. **(b)** Concordance matrix between the combined network (SNF) and each omics network based on NMI calculation (0=no mutual information, 1=perfect correlation). **(c)** SNF-combined similarity network colored by clusters (SNF-1/healthy-like=blue, SNF-2/severe at-risk=yellow, SNF-3/mild/moderate at-risk=grey) obtained after spectral clustering. Edges’ color indicates the strength of the similarity (black=strong, grey=weak). **(d)** PCA plot of samples based on fused network. Samples are colored by condition **(e)** Boxplots of DAG from untargeted lipid classes separated by groups. Significant stars are displayed for each comparison with *FDR <0.05, **FDR<0.01, ***FDR<0.001 (LIMMA). **(f)** Boxplots of TAG from untargeted lipid classes separated by groups. **(g)** PCA plot of samples after prior standardization based on significant metabolites between at least one pairwise comparison (LIMMA, FDR<0,05). Variance proportions are written on each component axis. Samples are colored by condition. **(h)** Circular heatmap of the top 159 metabolites (FDR<0.005). Metabolites are represented as slices and labeled around the plot. LogFC (LIMMA, FDR<0.05) from significant metabolites between groups are displayed in the first six outer layers. The 7^th^ to 9^th^ layers represent the coefficient of correlation between metabolites and BMI, metabolites respectively, and age (Spearman, p-value<0.1, R>0.15) and the p-value from significant associations between metabolites and gender (Chi-squared, p value<0.1). The inner layer represents the pathway of each metabolite.

### Lipids and metabolites highlight clinical differences between patient clusters

To characterize the molecular data-driven clusters of PLWH_ART_, we used clinical and single omics analysis. Cluster-specific clinical characteristics of PLWH_ART_ are presented in Table 1. Clusters were not statistically different for age, gender, duration of ART, and type of ART(p>0.05). On the other hand, SNF-1 had the healthiest profile (herein healthy-like group), SNF-3 an intermediate (herein mild/moderate group, and SNF-2 the most severe metabolic perturbations (herein severe at-risk group) indicating an at-risk metabolic profile. The severe at-risk groups represented patients with high BMI, central obesity, higher VAT, and incidence of MetS (all p<0.05). Regardless of disease severity, the severe at-risk group’s patients had a higher CD4+ T-cell count at the time of sample collection and more men who have sex with men (MSM) as transmission mode compared to the other clusters (all p<0.05). The at-risk groups, severe and mild/moderate had a significantly higher subcutaneous adipose tissue (SAT) and incidence of Hypertension compared to the healthy-like cluster (all p<0.05). The healthy-like cluster had the lowest BMI, SAT, VAT, and incidence of Hypertension (all p<0.05). A similar lipid class profile was observed between the healthy-like, mild/moderate at-risk groups, and HC (Table S2). Patients from the severe at-risk group showed a significant increase in diglycerides (DAG) (Fig 1E) and triglycerides (TAG) (Fig 1F) compared to healthy-like, mild/moderate at-risk cluster, and HC (all FDR<0.1) as well as other lipids classes which coordinate with their clinical metabolic profile (Fig S2). In this analysis, the relation between cluster and ART class was not significant (X2, FDR = 0,45). Still, we can mention that the three groups had an important proportion of missing data for this variable (16%, 29%, and 29% respectively). However, comparing the metabolites (Table S3), the mild/moderate at-risk group was the closest to HC, followed by the severe at-risk group, and finally, the healthy-like group, the centroid of which was close to the severe at-risk group centroid (Fig 1G). The differential metabolite abundance (DMA) was presented in Fig 1H. Among these highly differing metabolites, most perturbations were observed between HC and the healthy-like group (124/159) and HC and severe at-risk group (62/159). These clusters showed an up-regulation of the metabolites in the xenobiotics, nucleotides, and AA pathways compared to HC (HC vs. healthy-like group, 97/124, HC vs. severe at-risk group, 45/62). The mild/moderate at-risk group and HC had only nine metabolites differing, in line with the high clustering of both groups shown with PCA. In turn, the healthy-like and severe at-risk groups showed similar metabolic profiles. The lipids (mild/moderate vs. severe, 10/19, healthy-like vs. mild/moderate, 12/21) and amino acids (severe vs. mild/moderate, 7/19) differed significantly between the groups. Among these metabolites, 50 had a low or moderate association with age and BMI (Spearman correlation, absolute R<0.4, p<0.1) and 51 with gender (chi-squared test, p<0.1), showing the modest influence of individual characteristics on metabolomics profile. Combining the in-depth metabolomics and lipidomic data indicated more personalized risk factors for PLWH_ART_ that cannot be explained by the clinical features and a complex interplay between the multi-omics layers define overall health status.

### Sexual preferences influence the clusters’ differences driven by the microbiome

As the metabolic aberrations were closely linked with the microbiome profile, we investigated the microbiome’s impact in PLWH clusters. The α-diversity indices indicated a loss of diversity according to Observed, ACE, se.ACE, Chao1, and Fisher indices in healthy-like compared to the severe at-risk group (Mann Whitney, FDR<0.05) (Fig 2A, Fig S3, Table S4). A non-metric multidimensional scaling (NMDS) ordination of the dissimilarity-based index (Bray-Curtis) of diversity at the OTU level was performed to measure the inter-individual differences between groups (β-diversity) (Fig 2B). Based on NMDS plot axis coordinate 1, the healthy-like group was segregated separately from mild/moderate and severe at-risk groups (Mann Whitney, FDR<0.05, Fig 2C). The relative abundance of fecal microbiota was more influenced by the transmission mode than the cluster itself (Fig S4a). No other comorbidities on the microbiome profile were observed (Fig S4b-d). The severe at-risk group had a significantly higher number of MSM compared to the other groups (Table 1). While combining severe and mild/moderate at-risk groups, there were 69% (54/78) MSM in the at-risk clusters and 47% (9/19) MSM in the healthy-like group. This indicated that sexual preferences and the HIV-1 transmission mode relate to compositional differences in fecal microbiota between clusters. Permutational multivariate analysis of variance (PERMANOVA) at the family level showed that the centroids of the healthy-like groups were different from the severe at-risk (FDR<0.001) and mild/moderate groups (FDR=0.0054) (Table S5), indicating that there is only a location effect as permutation test for homogeneity of multivariate dispersions was not significant between the clusters (FDR>0.05). No statistical difference was observed between the severe and mild/moderate at-risk groups in both tests (FDR=0.38). The healthy-like group was enriched in Bacteroides and Lachnospira, while at-risk groups were enriched in Prevotella, Veillonella, and Succinivibrio (Fig 2D-2E). These families were also among 54 significantly discriminative features between healthy-like and at-risk groups as shown with linear discriminant analysis effect size (LefSe) (Fig 2F). Mann Whitney U test between clusters at the family level also found Prevotella and Bacteroides to be statistically distinct between these clusters (FDR<0.05, Table S6). Our data thus support the potential role of the Prevotella and Bacteroides in the cluster separation that could be mediated by the sexual preferences in PLWH_ART_.

**Figure 2:**
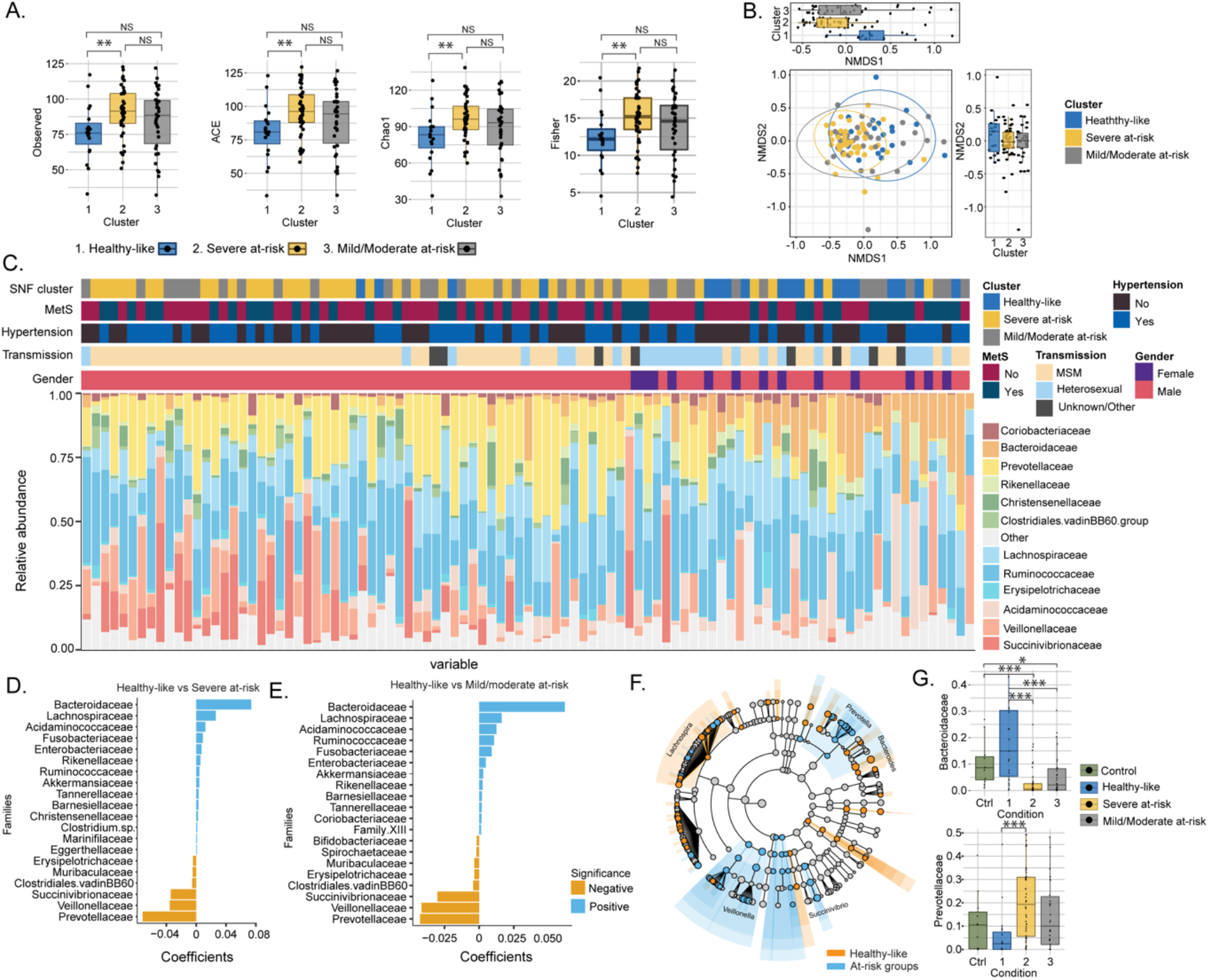
Transmission mode drove cluster differences in microbiome data. **(a)** Boxplots of alpha diversity indices (Observed, ACE, Chao1, Fisher) separated by HIV cluster. Significant stars are shown for each comparison (Mann-Whitney U test). **(b)** Non-metric multidimensional scaling (NMDS) plot of Bray-Curtis distances. Samples are colored by clusters. Boxplots based on NMDS1 and NMDS2 are represented. **(c)** Barplot represents the relative abundance of bacteria at the family level for each patient. Patient information is displayed above the barplot, including cluster, metabolic syndrome (MetS: yes/no), Hypertension (yes/no), transmission mode, and gender. **(d)** Barplot showing the top microbial families by representing their coefficient from PERMANOVA between SNF-1 and SNF-2. **(e)** Barplot showing the top microbial families between SNF-1 and SNF-3. **(f)** LEfSe cladogram representing cluster-specific microbial communities to healthy-like and to at-risk groups (SNF-2/SNF-3). Top families from PERMANOVA are labeled. **(g)** Boxplot of relative abundance at family level for Bacteroides (top) and Prevotella (bottom). Significant stars are shown for significant comparisons (Mann-Whitney U test).

### Factor and network analysis indicated the importance of microbiome-derived metabolites

To identify the molecular and clinical factors driving SNF cluster separation at the multi-omic level, we employed the Multi-Omic Factor Analysis (MOFA) tool for the multi-omics integration ^16^. After low variance filtering, the MOFA model was built using three views: microbiome with 173 OTUS, metabolome with 676 metabolites, and lipidome with 709 lipids. The model found 15 uncorrelated latent factors (Fig S5), i.e., combinations of features at the multi-omic level. The total variance was explained at 80% by the lipidome, 22% by the metabolome, and 2% by the microbiome, agreeing with the SNF analysis (Fig 3A). No factor explained most of the variance in the three views (Fig 3B). After selecting features with the largest weight in each cluster-associated factor (Fig 3C). Features with the most importance based on the top 10% of absolute weight were selected in each view, resulting in 396 features (263 lipids, 111 metabolites, and 22 OTUs). A good cluster separation based on hierarchical clustering of Spearman correlation confirmed the relevance of this subset of features (Fig 3D). We also extracted the top 20 features for each view based on this subset (Fig 3E). Bacteroides and Firmicutes were found in the phylum with the highest weight confirming our results from microbiome analysis and the importance of these microbial communities for cluster separation. Nevertheless, the microbiome had a lower weight than metabolites and lipids in MOFA factors. Among the top 20 metabolite features, three metabolites derived or partially derived from microbiota (MDM) (3,4−dihydroxybutyrate, 2−oxindole−3−acetate, and indoleacetylglutamine) were found (Fig 3E). To investigate the coordinated role of MDM, we performed the consensus association analysis (Fig S6). To balance the different number of features in each of the three omics, we randomly selected 241 metabolites, 241 lipids, and 241 OTUs 1000 times. Significant pairwise correlations (FDR<10^−6^) found in >90% of comparisons were used to build a positive co-expression network, and community detection was performed, resulting in a network with 1324 nodes, (694 lipids, 536 metabolites, 94 microbial communities), 131863 edges and eight multi-omic communities (N > 30) (Fig S11). To refine this network, we selected the 396 features based on MOFA differing the most clusters (Fig 3D) in the co-expression network (Fig 3F). The most central communities (Average degree C1=444, Average degree C2=364) were lipid specific (SNF-1, lipids=122/124, SNF-2, lipids=127/128), while metabolites enriched communities were sparser with a lower average degree (C3=26, C4=22, C6=10, C7=6) but still connected to lipids with 86 edges between lipids and metabolites. Microbiome enriched community (c8) did not correlate with metabolites or lipids. However, eight MDMs were found in the network, mostly in c6 (5/21), showing that MDMs were highly intercorrelated and can have a potential role in shaping the systemic metabolic and lipid profile.

**Figure 3:**
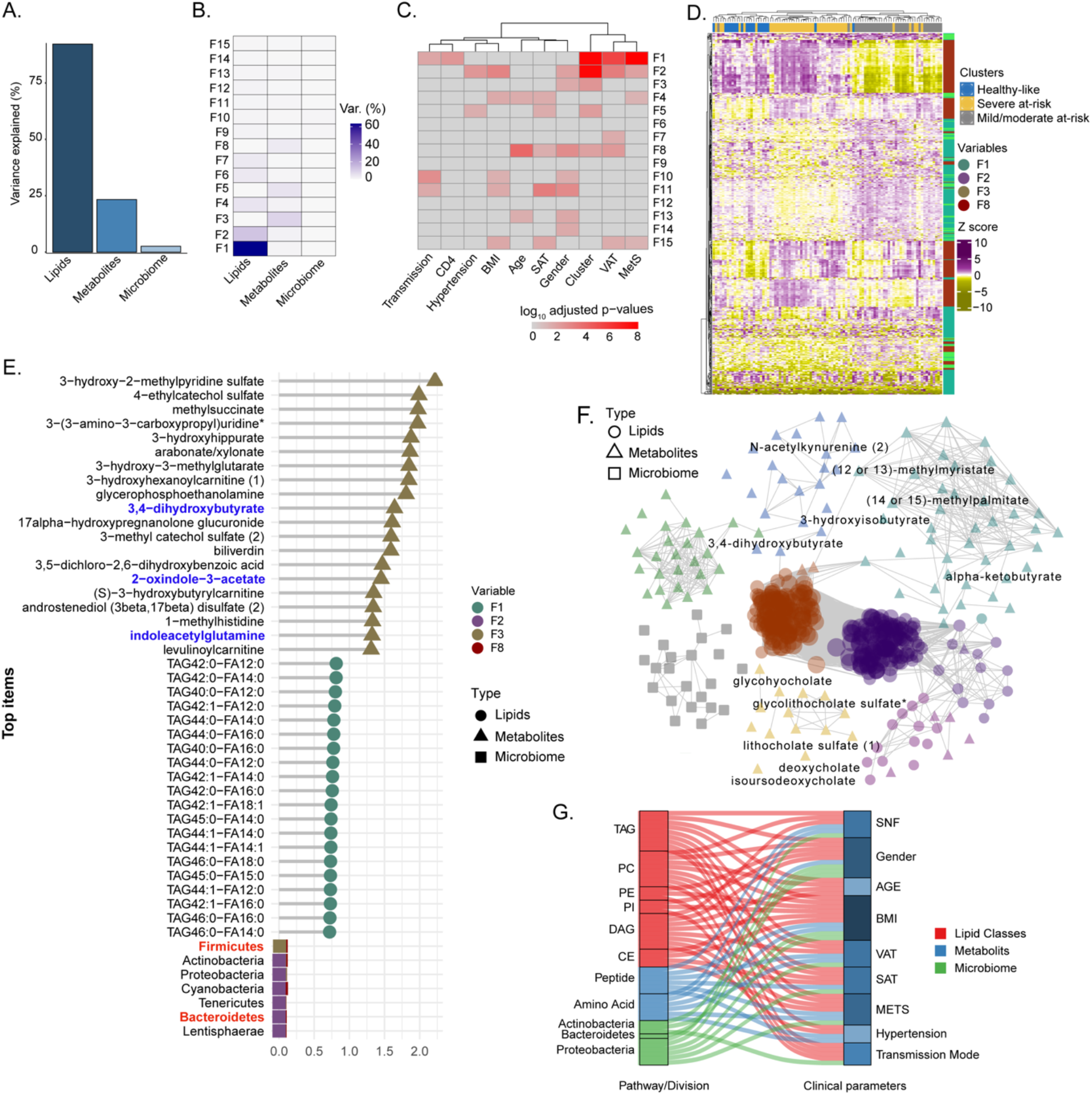
Factor analysis highlights the essential features for cluster separation and potential microbiome-derived metabolites importance. **(a)** Barplot of total variance explained by MOFA model per view. **(b)** Variance decomposition plot. The percentage of variance is explained by each factor for each view. **(c)** External covariate association with factors plot. Association is represented with log10 adjusted p-values from Pearson correlation. **(d)** Heatmap representing levels of microbial communities, metabolites, and lipids with the higher absolute weight in MOFA factors associated with cluster (F1, F2, F3, F5, F8). Samples are labeled according to the study groups. Data were Z-score transformed. The type of data (lipid, metabolite, microbe) is displayed on the right. **(e)** Top 20 features with higher absolute weight in MOFA factors associated with cluster (F1, F2, F3, F5, F8) from lipidome, metabolome, and microbiome. Microbiome-derived metabolites and bacterial phylum of interest are colored in blue and red, respectively. **(f)** MOFA features differing clusters and interactions extracted from the 3-layers consensus co-expression network. Microbiome-derived metabolites are labeled.

### MDM is highly associated with clinical features driven by bile acid metabolism and indole derivatives

We observed a high correlation among the MDMs (Fig 3F). Therefore, to further investigate their the role in PLWH, we retrieved 69 metabolites defined as, i) produced by intestinal bacterial mainly part of secondary bile acid metabolism (n=22) and ii) produced by host modified by bacteria (n=47, polyamines, propionate, acetate, butyrate, and indole derivatives) as reported (Table S7) ^17^. Differential abundance analysis 19 MDMs differed between HC and PLWH irrespective of the SNF clusters (Fig 4A). The propionate and indole derivates were significantly (FDR*<0*.*05*) increased in PLWH compared to HC. As observed in the whole metabolomics profile, mild/moderate had a more similar profile to HC than healthy-like and severe at-risk groups, while healthy-like and severe at-risk group had identical profiles. We performed univariate linear regression to investigate the link between microbiome-derived metabolites and clinical parameters (Table S8). Lithocholate sulfate was associated with obesity-related comorbidities (MetS, SAT, VAT, Hypertension, central obesity) and 2-aminobutyrate and deoxycholic acid 12-sulfate. Several lifestyle parameters impacted MDM, such as poultry and vegetable intake, smoking, and alcohol. The use of medication as antihypertensives was also associated with three MDMs. Glycolithocholate and glycoursodeoxycholic acid sulfate were linked to HIV-related parameters (CD4 nadir, CD4 at study entry) and patients’ demography and lifestyle parameters. The SNF cluster was linked to lithocholate sulfate, 3-ureidopropionate, and imidazole propionate (Fig 4B). Finally, to measure the influence of MDM on plasma metabolomics profile, we performed association analysis and community detection on metabolomics data only (Fig 4C). We obtained a co-expression network with 843 nodes and 15490 edges (FDR<0.02) and observed seven communities (c1-c7) (Fig 4C). The c4 contained all the secondary bile acid metabolites. Though the differential abundance analysis did not show all MDM differences between the SNF clusters and HC, they were highly correlated in PLWH, with significant MDMs differing between the groups (Fig 4D). Combining all the data, we showed an essential role of MDMs in the system-level metabolic profile of PLWH on successful therapy.

**Figure 4:**
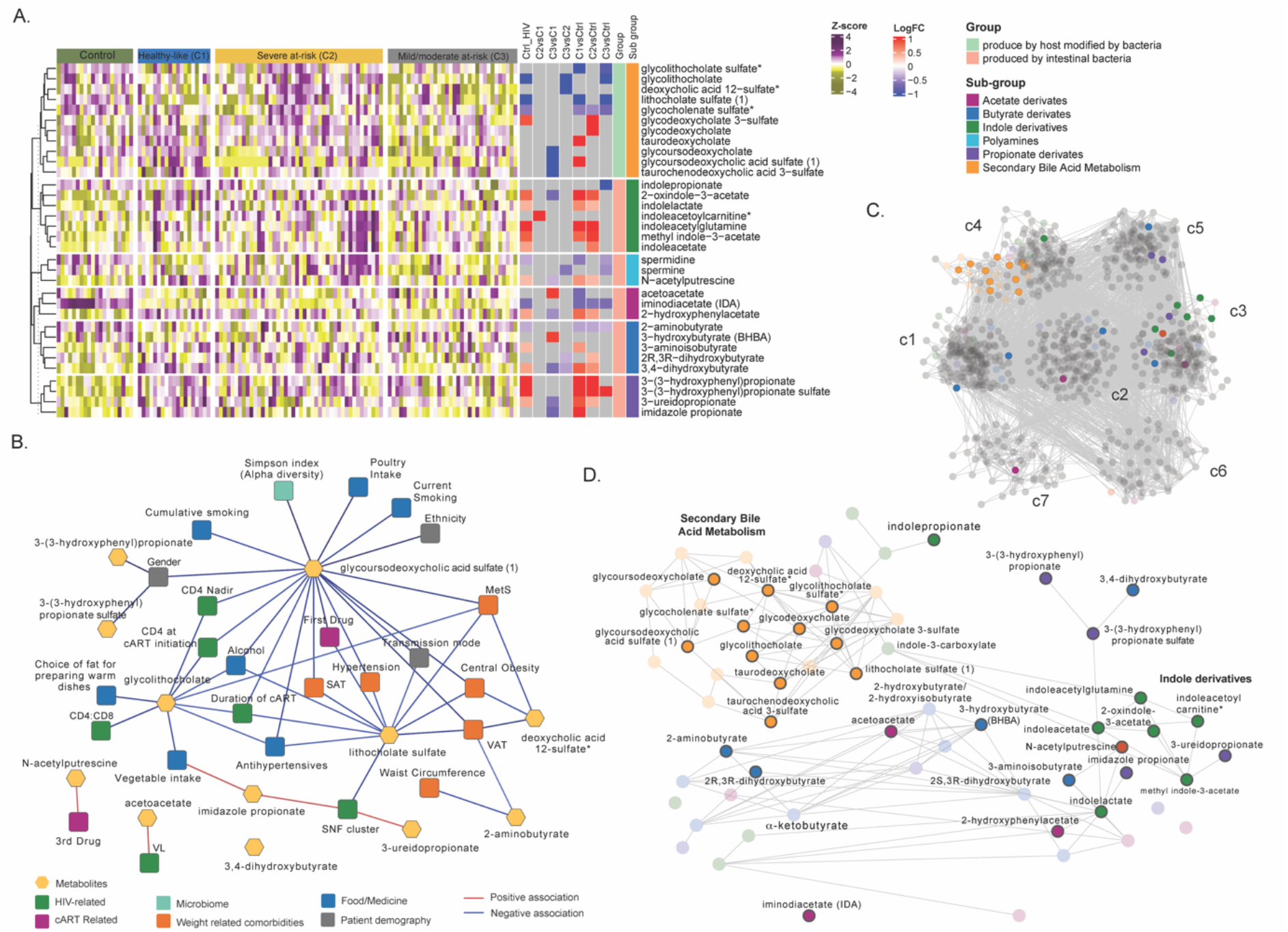
Microbiome-derived metabolites are affected in HIV clusters. **(a)** Heatmap representing abundances of microbiome-derived metabolites differing at least one comparison. Data were Z-score transformed. Significant logFC (LIMMA, FDR < 0.05) of pairwise comparisons between conditions, groups, and under groups of microbiome-derived metabolites are displayed on the right. **(b)** Cytoscape network showing significant positive and negative associations between clinical parameters and microbiome-derived metabolites (univariate linear regression, FDR<0.05). Clinical parameters are colored based on categories. **(c)** Co-expression network of metabolomics data in PLWH. Metabolites are grouped by communities, and microbiome-derived metabolites are labeled and colored based on the subgroup. **(d)** The subset of microbiome-derived metabolites from the co-expression network. Non-significant metabolites in all comparisons are displayed with transparency. Significant microbiome- derived metabolites between at least two conditions are labeled.

## Discussion

In this study, we used network and factorization-based integrative analysis of plasma metabolomics, lipidomics, and microbiome profile to characterize clinical phenotypes in the PLWH_ART._ We identified three different diseases’ state -omics phenotypes (SNF-1 to SNF-3) within PLWH_ART_ driven by metabolomics, lipidomics, and microbiome that a single omics or clinical feature could not explain. The integrative omics highlighted the importance of highly intercorrelated microbiome-derived metabolites and their association with the clinical parameters in PLWH_ART_ clusters separation shaping their systemic health profile. The severe at-risk group (SNF-2) has the at-risk metabolic profile characterized by an increase in TAG and DAG, highest median BMI, MetS incidence, VAT, and SAT, but had a higher CD4 T-cell count at sample collection compared to healthy-like and mild/moderate at-risk group, which displayed an HC like lipidomic profile. However, the healthy-like and severe at-risk group had a similar metabolic profile differing from HC, with dysregulation of AA metabolism. At the microbiome profile, the healthy-like group had a lower α-diversity, a lower proportion of MSM, and was enriched in Bacteroides. In contrast, in at-risk groups, there was an increase in Prevotella, with a high proportion of MSM confirming the influence of sexual orientation on the microbiome profile ^8^. Our study thus identified a risk group of PLWH with successful treatment with a dysregulated metabolic profile potentiate metabolic diseases that could be barriers to healthy aging.

Similarity network analysis reduces the high-dimensional nature and different variance of multi-omics data to group patients based on the most similar profile ^15^. One of the main advantages of this method is the possibility to compare the networks’ similarities to find out which layer has the most similarity with the final network. The similarity network fusion-based patient stratification has been used primarily in non-communicable diseases like cancer [to identify cancer subtypes ^15, 18^ and prognosis ^19^], respiratory diseases ^20^ and to study the influence of diet on human health ^21^. Recently we developed SNF-based patient stratification by integrating transcriptomics and metabolomics to define disease severity in COVID-19 that are predictive of the most robust biological features ^22^. We also reported the influence of gut microbiota on the systemic metabolic profile associated with disease severity ^23^. However, no data were presented to stratify the PLWH_ART_ to fingerprint their disease status. The SNF has shown that the most crucial omics layer in cluster separation was lipids (NMI=0.6), supported by the MOFA analysis. A study reported that ART and HIV reservoirs are responsible for changes in adipose tissue and lipids metabolism in PLWH ^24^. Dyslipidemia represents the increase in triglycerides, low-density lipoprotein cholesterol (LDL-C), total cholesterol (TC), and decrease of high-density lipoprotein cholesterol (HDL-C) cholesterol in the blood is a well-recognized complication observed in PLWH both naïve ^25^ and after ART initiation leading to cardiovascular diseases and mortality ^26, 27^. We found that the severe at-risk individuals (44/97) had most lipids classes upregulated, especially TAG, DAG, and CER, compared to the other groups, while healthy-like and mild/moderate at-risk groups had no difference with HC. The severe at-risk group also has more patients with high BMI, VAT, SAT, and incidence of MetS. DAG and TAG high levels have been linked to cardiovascular events ^26, 28^. The TAG levels have been linked to insulin resistance and increased diabetes risk ^26^, confirming this cluster group’s qualification as patients with dysregulated lipid profiles and metabolic disease risk. The association of lipid profile with CD4 counts is still debated. It is positively associated with ^27, 29^, and negatively ^30^ associated with the high abundant lipid profile. Interestingly, we found the severe at-risk group to have the highest CD4 count and suppressed viremia but have dysregulated lipid profile that could be reasoned for unhealthy aging and adverse cardio-metabolic health. Therefore, we propose using a holistic view to define the clinical and immunological treatment success of PLWH_ART_ beyond viral suppression and immune reconstitution.

The second omics defining clusters were metabolites (NMI=0.4). Interestingly, the metabolic profile was not completely overlapping with the lipid profile showing the complexity associated with the disease. PLWH_ART_ in the healthy-like group were the most different from the HC regarding their healthy-like clinical parameter with the lowest BMI, VAT, and SAT. Nevertheless, 32% of PLWH_ART_ in the healthy-like group had MetS, which was half of the severe at-risk group (70%) but double the mild/moderate at-risk group (17%), indicating a possible lipid-independent metabolic dysregulation. Still, the mild/moderate at-risk group had the profile of the most HC-like, similarly to the lipids, despite having a significantly higher number of patients with Hypertension than the healthy-like group. The healthy-like and severe at-risk groups showed an up-regulation of the metabolites in the xenobiotics, nucleotides, and AA metabolism, indicating a potential role of diet. We previously showed that the glutamate metabolism was highly disrupted in PLWH_ART_ with MetS in the same COCOMO cohort ^11^, which can be responsible for late immune recovery in the short-term ART patients ^31^. Also, short-chain dicarboxylacylcarnitines (SCDA) and glutamine/valine were higher in PLWH with coronary artery disease than in controls ^32^. In our cohort, we observed glutamate, N-acetyl-glutamate, phenyl-acetyl-glutamate, gamma-glutamylglutamate, and 4-hydroxyglutamate to be upregulated between severe at-risk and healthy-like groups than HC and mild/moderate at-risk group. 4-hydroxyglutamate was increased in the severe at-risk group compared to the healthy-like group.

The microbiome network had a modest similarity with the final SNF network (NMI=0.3), and clustering was not observed on the PCA plot. Metabolism and immunity of the host have been shown to be affected by bacteria and disrupted microbiome linked to illness ^33^. More importantly, there is a high variability of microbiota among individuals based on lifestyle, diet, medication, and physiology ^34^. Increased α-diversity is associated with good health and decreased diversity in several diseases, including HIV ^6^. A meta-analysis reported that HIV status was not associated with decreased a-diversity in MSM, perhaps due to sexual behaviors, but was decreased in PLWH with heterosexual transmission ^35^. Despite having healthy clinical and metabolic profiles, we observed an increase in α-diversity in the severe at-risk group compared to the healthy-like group driven by MSM and no differences between the healthy-like and mild/moderate at-risk group. In terms of bacterial composition, studies reported that PLWH has a higher abundance of Prevotella and a lower abundance of Bacteroides^36^. The crucial point is that the ratio is associated more with MSM than HIV status and has been shown by several studies ^6-8^. Our study observed that the severe at-risk group was enriched in Prevotella and depleted in Bacteriodes compared to the healthy-like group. Interestingly, the decrease of Bacteroides in obese patients was inversely correlated with serum glutamate ^37^, which was also observed in severe at-risk group patients. On the other hand, some Prevotella species have proinflammatory effects, leading to intestinal inflammation, bacterial translocation, and microbiome dysbiosis ^38^. In general, the cohort is mainly composed of MSM (63/97). As described above, it confirmed that the difference in the microbiome is driven by MSM status in severe at-risk groups, as the patients have 81% MSM. The mild/moderate at-risk group, even if there is no difference from the severe at-risk group according to PERMANOVA, has the same proportion of MSM as the healthy-like group. It has been proposed that early regulation of the MSM-related microbiome could help prevent HIV infection ^6^. However, the question remains whether the MSM-related microbiome is a potential driving force of metabolic comorbidities or whether MSM is a confounding factor disturbing a potentially clinical signal from a disturbed microbiome.

Microbial compositions have implications for metabolism and metabolic diseases, notably through the production of MDMs ^39^. Secondary bile acids transformed from primary bile acids by bacteria have a role in lipid digestion. It regulates host metabolism through signaling and can inhibit the production of proinflammatory cytokines by immune cells ^17^. Lipid metabolism, including triglyceride trafficking, is influenced by bile acids through the interaction with the Farnesoid X receptor (FXR) receptor and has been implicated in mice’s metabolic disorder ^40^. A bile acid, glycolithocholate was found upregulated in healthy-like and mild/moderate at risk-group compared to controls shown previously associated with insulin resistance ^41^. It was highly negatively associated with food elements such as vegetable intake and choice of fat for cooking, alcohol, and HIV-related parameters such as CD4 levels (nadir and at ART initiation) and HIV duration. High glycodeoxycholate was observed in healthy-like especially compared to mild/moderate at-risk group, while the glycodeoxycholic acid has been shown to be negatively associated with the insulin resistance ^37^. Glycocholenate sulfate was down-regulated in the three clusters compared to controls. All secondary bile acids were shown to be highly intercorrelated in co-expression analysis. Three other bile acids, lithocholate sulfate, glycousodesoxycholic acid sulfate, and deoxycholic acid 12-sulfate, were negatively associated with metabolic perturbations including MetS, VAT, and central obesity. Acetate, propionates, and butyrate are part of short-chain fatty acids (SCFAs) and are obtained from the fiber bacterial fermentation in the colon that the host’s enzymes cannot digest ^42^. Proprionate derivates were upregulated in healthy-like and severe at-risk groups. Acetate and butyrate derivates had a more variable profile. Imidazole propionate (IMP) and 3-ureidopropionate were linked to the SNF clusters. In our study, the IMP was also linked to vegetable intake, which was reported to be involved in the insulin resistance ^39^. The Bacteroides metabolize most of the acetate and propionate from polysaccharides, and Firmicutes produce butyrate ^17^, which does not explain the relationship within the SNF clusters indicating a more complex interplay between the MDMs and bacterial community in a diseased condition. Tryptophan is converted by bacterial tryptophanase into indole, and indole derivates are involved in the host-microbiota homeostasis ^43^. Indoles derivates mainly were upregulated in the healthy-like and severe at-risk groups. Our data thus suggested the role of MDMs in shaping the clinical phenotype and systemic health profile in PLWH_ART,_ which could be a therapeutic target for improving health.

Though our study is the first to demonstrate an integrative multi-omics approach to the role of MDMs in systemic alterations in PLWH_ART_, our study has limitations that merit comments. First, the study is cross-sectional and therefore restricted to predicting dynamic interactions of different omics layers. Second, the microbiome data analysis is through 16S methodologies with a higher level of missing data at the genus and species level than metagenomics. Third, although the network-based analysis and the observational data suggest a potential causal association of altered metabolic profile with clinical features, other factors may drive observed effects. Fourth, though this is the largest study to date to perform integrative omics in PLWH, the number of samples was relatively low. Finally, both microbiome and metabolomics are highly dependent upon the genetics, environment, and diet of an individual. The interaction noted may characterize the epiphenomena of a personalized immune system that can be an avenue for future studies to develop a more personalized model for integrative omics to phenotype the disease states that we recently reported ^22^.

In conclusion, we performed a multi-omics analysis of PLWH_ART_ with different clinical features. We identified the diversity of PLWH_ART_ in HIV-related biological alterations regardless of immunological recovery and virological suppression. A proportion of PLWH_ART_ (SNF-2; severe at-risk group around 45% in the present cohort) showed highly dysregulated lipidomics (increased TAG and DAG) and clinical profile (increased BMI and obesity-related features) with increased Prevotella and decreased Bacteroides, the latter being related to MSM transmission. However, alterations in the metabolomics profile and higher CD4 T-cell count at the time of sample collection indicate a complex systemic interplay between host immunity and metabolic health that might affect healthy aging in this population. Integrative analytical approaches that reflect the overall systemic health profile of PLWH_ART_ may improve patient stratification and individual therapeutic and preventive strategies. Developing a more personalized model or targeting the interaction networks rather than individual clinical or omics features may provide novel treatment strategies in countering dysregulated metabolic traits, aiming to achieve healthier aging.

## Materials and Methods

### Patient Cohort and Multiomics data

The cohort comprises 97 PLWH_ART_ from the Copenhagen Comorbidity (COCOMO) Cohort, a prospective cohort of PLWH_ART_. This study used an untargeted metabolomics ^11^, a complex lipid profile ^13^, and a 16s microbiome data ^7^ reported earlier for the larger cohorts. We also extracted clinical and demographical data from the COCOMO database. The HIV-negative controls (HC) (n=20) were used to understand the basal level of omics.

### Similarity network fusion (SNF)

Lipids and metabolites with low variance (<0.3) were removed from the data. The three layers of omics (microbiome, lipidome, metabolome) were standard normalized before analysis. Analysis was processed using the package SNFtool as described ^15^. Pairwise sample distances were calculated with the function dist2 followed by the construction of similarity graphs (number of neighbors, K=13, hyperparameter, alpha=0.8) for each layer. The similarity network fusion (SNF) was used to fuse all the networks (K=13, number of iterations, T=10). Spectral clustering was applied to the fused network (Number of clusters C=3). The parameters (K, alpha, T, C) were chosen to maximize the Eigen gap and minimize rotation cost. A concordance matrix was calculated based on the similarity in cluster assignments in each network, and the fused network was calculated in normalized mutual information (NMI).

### Clinical data statistics

Clinical characteristics were compared between clusters using pairwise tests. Welch’s T-test and Mann-Whitney U tests were used to compare normally distributed and non-normally distributed continuous variables were compared using R ^44^. Chi-Square Test was used to compare discrete variables if the expected values of the contingency table were five or more. Otherwise, Fisher’s Exact Test was used. Univariate linear regression was performed with the function lm, and correlations were calculated using the cor.test and cor functions from the R stats package ^44^.

### Lipidomics and metabolomics analysis

Untargeted metabolomics and lipidomics were log2 transformed before analysis was performed in Metabolon™, USA. Lipid data were grouped by lipid classes as in the following.

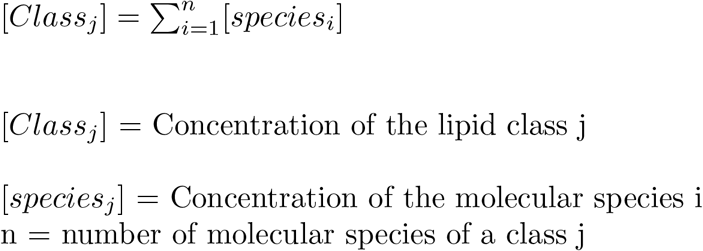

Differential abundance analysis was performed pairwise with the R package limma between groups (HC, SNF-1, SNF-2, SNF-3) for each data set (lipidomics classes, lipidomics, metabolomics). Benjamini-Hochberg (BH) adjustment was applied, and FDR was set up to 0.05 for statistical significance. Deviations were mentioned in the respective analysis. The default p-value cutoff was set to 0.05. Other p-values cutoffs are adapted for a specific analysis depending upon the number of significance and to minimize the false positivity.

### Microbiome analysis

Microbiome data analysis was performed using the R package phyloseq ^45^. Alpha diversity estimates were calculated using the estimate_richness function and the following measures: Observed, ACE, se.ACE, Chao1, Shannon, Simpson, InvSimpson, and Fisher. NMDS ordinations based on Bray-Curtis distances between all samples were calculated using the ordinate function. Otu table was converted to relative abundances for further analysis. The vegan package ^46^ was used to perform PERMANOVA. Equal multivariate dispersion was verified using the betadisper function applying Marti Anderson’s PERMDISP2 procedure. Pairwise PERMANOVA test was done between groups using the adonis function, Bray distance, and Bonferroni correction. The cutoff for the adjusted p-value was set up to 0.05. Galaxy module LDA Effect Size (LEfSe) was used to find microbial communities (at genus, family, or higher level) specific to one specific cluster ^47^. The multiclass analysis approach was one against all. First, a non-parametric factorial Kruskal-Wallis (KW) sum-rank test was performed with clusters (cutoff alpha=0.05), followed by pairwise Wilcoxon rank-sum tests between clusters (cutoff alpha=0.05), and then effect size calculation for each significant feature was done using discriminant analysis (absolute LDA score>2). Results are represented using a cladogram produced by the module.

### Microbiome-derived metabolites

Microbiome-derived metabolites, groups, and subgroups were retrieved from the previous literature ^17^.

### Multi-omics Factor analysis (MOFA)

Filtered data for SNF was also used for MOFA analysis ^16^. Microbiome data were rarefied by filtering based on variance (>0.2). In addition, the microbiome data were center log-ratio (CLR) transformed to follow a normal distribution. The MOFA model was trained using default parameters, and sample metadata was added to the model. Total variance explained per view was used to see the weight of each omics layer. A correlation plot was used to verify the low correlation between factors. A variance decomposition plot was used to determine the percentage of variance explained by each factor and omics layer. Association analysis of the factors with clinical features was done using MOFA function correlate_factors_with_covariates and factors associated with the SNF cluster selected. 5 and 95 % quantile weights for each view were selected for each factor. Pathway analysis was performed on factors using the MOFA function run_enrichment for each view, with the parametric statistical test, FDR-adjusted p-values, and separated positive and negative values. Annotation libraries were made from Metabolon™ super pathways for metabolomics and lipidomics and Division level for the microbiome.

### Co-expression analysis

Pairwise Spearman correlations between features were calculated, and the cutoff for FDR of significant correlations was selected to minimize the number of false positives. The positive and negative networks were built using the python igraph ^48^ and compared to random networks of the same size. Leiden community detection was applied to find groups of interconnected features, and the mean degree was calculated to represent the community centrality using python module leidenag ^49^. Communities of less than 30 features were excluded. Consensus association analysis was performed to integrate the three layers of omics using 1000 iterations. At each iteration, pairwise correlations between OTUs (N=241), 241 metabolites, and 241 lipids selected randomly were run, and significant positive correlations (Spearman, FDR<0.001) were kept as an association. Associations found in 90% of the comparisons over all iterations were kept building the final network as described above.

### Visualization

Scatter plots, PCA plots, box plots, NMDS plots, circular heatmap, and bar plots were generated using ggplot2 ^50^. Heatmaps were generated using ComplexHeatmap ^51^. Sankey plot was made using the R package ggalluvial ^52^. Networks were plotted using Cytoscape v3.6.1 ^53^.

## Data and Code Availability

All of the data generated or analyzed during this study are included in this published article and/or the supplementary materials. Created datasets and code are publicly available. The metabolomics and lipidomics data are available from 10.6084/m9.figshare.14356754 and 10.6084/m9.figshare.14509452. All the codes are available at github: https://github.com/neogilab/HIV_multiomics

## Ethical clearances

Ethical approval was obtained by the Regional Ethics Committee of Copenhagen (COCOMO: H-15017350) and Etikprövningsmyndigheten, Sweden (Dnr: 2022-01353-01). Informed consent was obtained from all participants and delinked before analysis.

## ACKNOWLEDGMENTS

The study is funded by the Swedish Research Council grants 2017-01330, 2018-06156, and 2021-01756 to UN. Novo Nordic Foundation, Lundbeck Foundation, Augustinus Foundation, Region Hovedstaden, and Rigshospitalet Research Council to SDN.

## Supplementary Tables

Table S1. List of parameters used in the study.

Table S2a. Table of differential lipid abundance analysis on individual lipids abundances.

Table S2b. Table of differential lipid abundance analysis grouped by lipids classes.

Table S3. Table of differential metabolite abundance analysis.

Table S4. Alpha diversity indices statistics.

Table S5. Permutational multivariate analysis of variance at the family level.

Table S6. Microbiome statistics at a family level.

Table S7. List of microbiome-derived metabolites

Table S8. Univariate linear regression between clinical parameters and microbiome-derived metabolites differing groups.

## Supplementary Figures

**Fig S1**. PCA plot of samples after prior standardization based on a) Lipidomics b) Metabolomics c) Microbiome. Variance proportions are written on each component axis. Samples are colored by condition (Ctrl = green, SNF-1 = blue, SNF-2 = yellow, SNF-3 = grey).

**Fig S2**. Boxplots of untargeted lipid classes are separated by groups. Color is based on groups (Ctrl = green, SNF-1 = blue, SNF-2 = yellow, SNF-3 = grey). P values are displayed for each comparison (Mann Withney U Test).

**Fig S3**. Boxplots of alpha diversity indices (se.chao1,Simpson, Shannon, se.ACE, InvSimpson) separated by HIV-cluster. Color is based on groups (Ctrl = green, SNF-1 = blue, SNF-2 = yellow, SNF-3 = grey).

**Fig S4**. Non-metric multidimensional scaling (NMDS) plot of Bray-Curtis distances. Samples are colored by A) Transmission mode B) Central obesity C) Metabolic Syndrome D) Hypertension.

**Fig S5**. Correlation matrix of MOFA factors. Size and transparency are proportional to the absolute coefficient of correlation. Color is displayed as a gradient-based coefficient of correlation from -1 (red) to 1 (blue).

**Fig S6**. Cytoscape consensus co-expression network. Color and label are based on communities.

