## Supplementary material for "Network-based multi-omics integration reveals metabolic at-risk profile within treated HIV-infection": Table 1

**Table 1.** Patient characteristics

|  | Complete Cohort | SNF-1 | SNF- 2 | SNF-3 | P values |
| --- | --- | --- | --- | --- | --- |
| N | 97 | 19 | 44 | 34 |  |
| Age in years, Median (IQR) | 54 (48-63) | 60 (48-68) | 54 (48-62) | 54 (51-60) | 0.75 |
| Gender, Male, N (%) | 84 (87) | 15 (79) | 40 (91) | 29 (85) | 0.36 |
| Ethnicity Caucasian, N (%) | 79 (81) | 15 (79) | 38 (87) | 26 (77) | 0.49 |
| Mode of transmission, N (%)  Homosexual/bisexual  Heterosexual  Other/unknown | 63 (65)  26 (27)  8 (8) | 9 (47)  7 (37)  3 (16) | 36 (81)  6 (14)  2 (5) | 18 (53)  13 (38)  3 (9) | 0.017 |
| CD4 Nadir, cells/μL, Median (IQR) | 235 (123-320) | 240 (127-330) | 240 (145-365) | 223 (42-290) | 0.49 |
| CD4 at ART Initiation, cells/μL, Median (IQR) | 287 (155-410) | 270 (120-360) | 318 (192-463) | 240 (108-320) | 0.11 |
| Viral Load at ART initiation, log copies/mL, Median (IQR) | 5.02 (4.34-5.61) | 4.87 (4.32-5.5) | 5.11 (4.74-5.61) | 4.94 (4.2-5.55) | 0.35 |
| CD4 at sampling, cells/μL, Median(IQR) | 713 (570-900) | 680 (540-958) | 762 (689-923) | 610 (475-819) | 0.015 |
| CD8 at sampling, cells/μL, Median (IQR) | 775 (600-1100) | 780 (630-879) | 894 (638-1300) | 700 (530-870) | 0.054 |
| Viral load (<50 copies/mL), N (%) | 97 (100) | 19 (100) | 44 (100) | 34 (100) | 1 |
| Duration of treatment in years, median (IQR) | 15 (9-18) | 15 (13-18) | 15 (8 -18) | 14 (7-17) | 0.73 |
| Current Treatment, 1^st^ drug, N (%)  ABC  TDF/TAF  Other | 31 (32)  42 (43)  24 (25) | 8 (42)  8 (42)  3 (16) | 13 (30)  19 (43)  12 (27) | 10 (29)  15 (44)  9 (27) | 0.84 |
| Current Treatment, 3^rd^ drug, N (%)  NNRTI  PI/r  INSTI  Other | 38 (39)  18 (19)  15 (15)  26 (27) | 8 (42)  4 (21)  4 (21)  3 (16) | 14 (32)  11 (25)  6 (14)  13 (29) | 16 (47)  3 (9)  5 (15)  10 (29) | 0.45 |
| BMI, Mean (SD) | 24 (22-27) | 22 (19-25) | 26 (23-28) | 24 (22-27) | 0.003 |
| VAT, Median (IQR) | 89 (36-142) | 41 (19-106) | 127 (79-196) | 69 (26-100) | 0.0001 |
| SAT, Median (IQR) | 111 (70-167) | 69 (33-115) | 117 (82 -174) | 119 (83-190) | 0.02 |
| MetS, N (%) | 43 (44) | 6 (32) | 31 (70) | 6 (17) | 0.000009 |
| Hypertension, N (%) | 49 (51) | 5 (26) | 23 (52) | 21 (62) | 0.04 |
| Central obesity, N(%) | 57 (59) | 8 (42) | 32 (73) | 17 (50) | 0.033 |
| Waist circumference (cm) | 94 (87-101) | 90 (84-95) | 100 (91-105) | 90 (87-97) | 0.0007 |
