## Supplementary figures for "Network-based multi-omics integration reveals metabolic at-risk profile within treated HIV-infection"

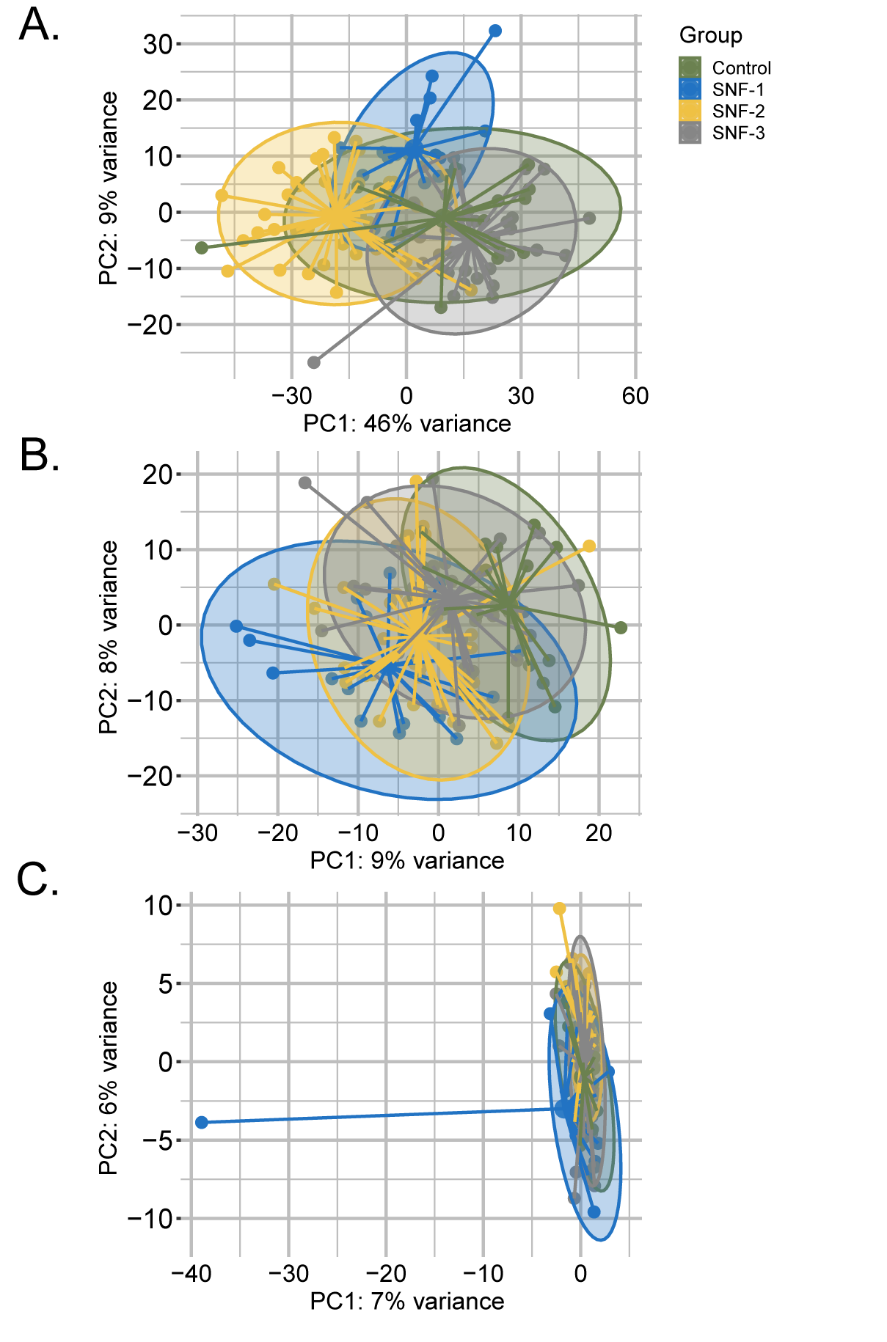

**Fig S1.** PCA plot of samples after prior standardization based on a) Lipidomics b) Metabolomics c) Microbiome. Variance proportions are written on each component axis. Samples are colored by condition (Ctrl = green, SNF-1 = blue, SNF-2 = yellow, SNF-3 = grey).

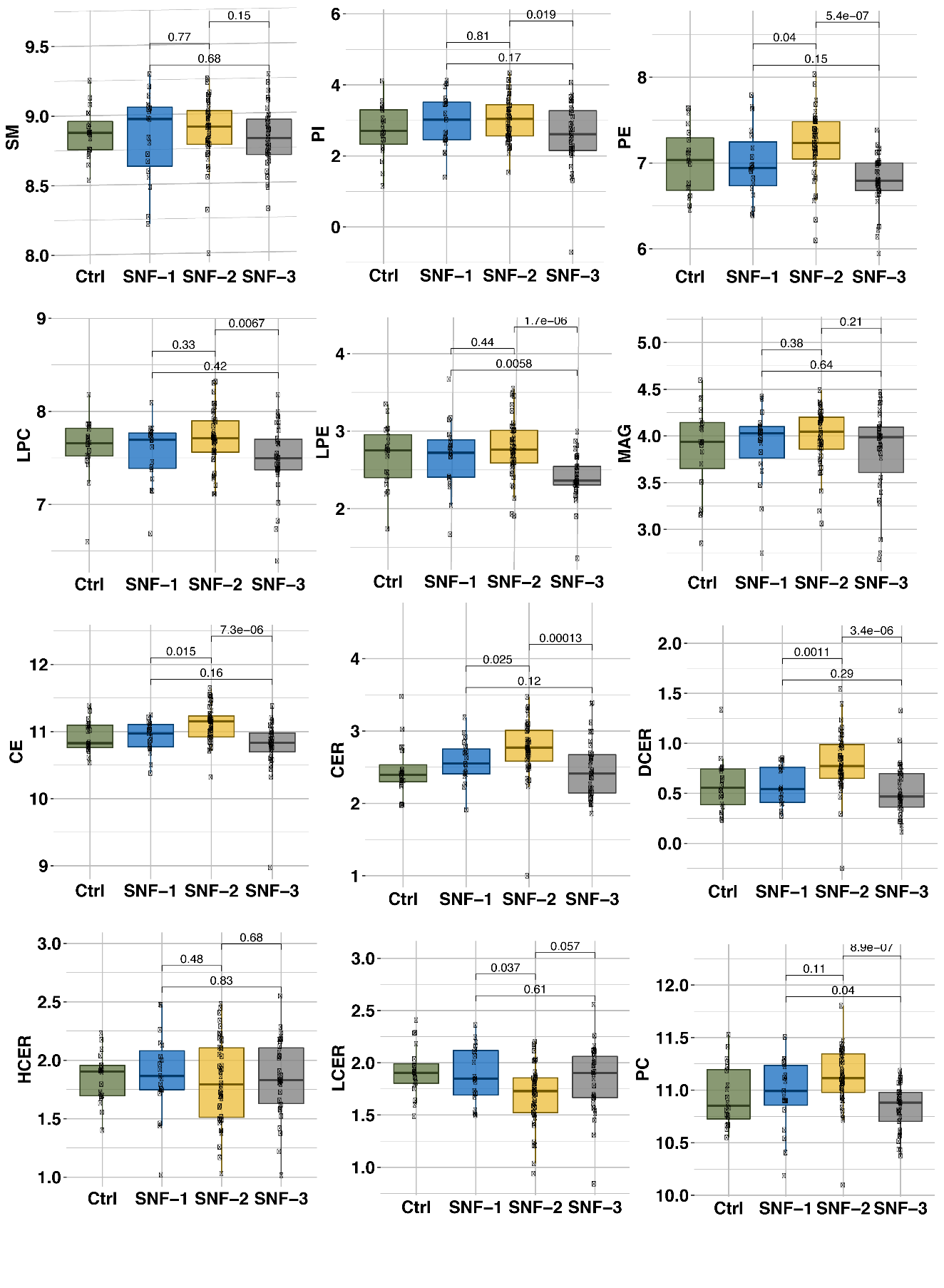

**Fig S2.** Boxplots of untargeted lipid classes separated by groups. Color is based on groups (Ctrl = green, SNF-1 = blue, SNF-2 = yellow, SNF-3 = grey). P values are displayed for each comparison (Mann Withney U Test).

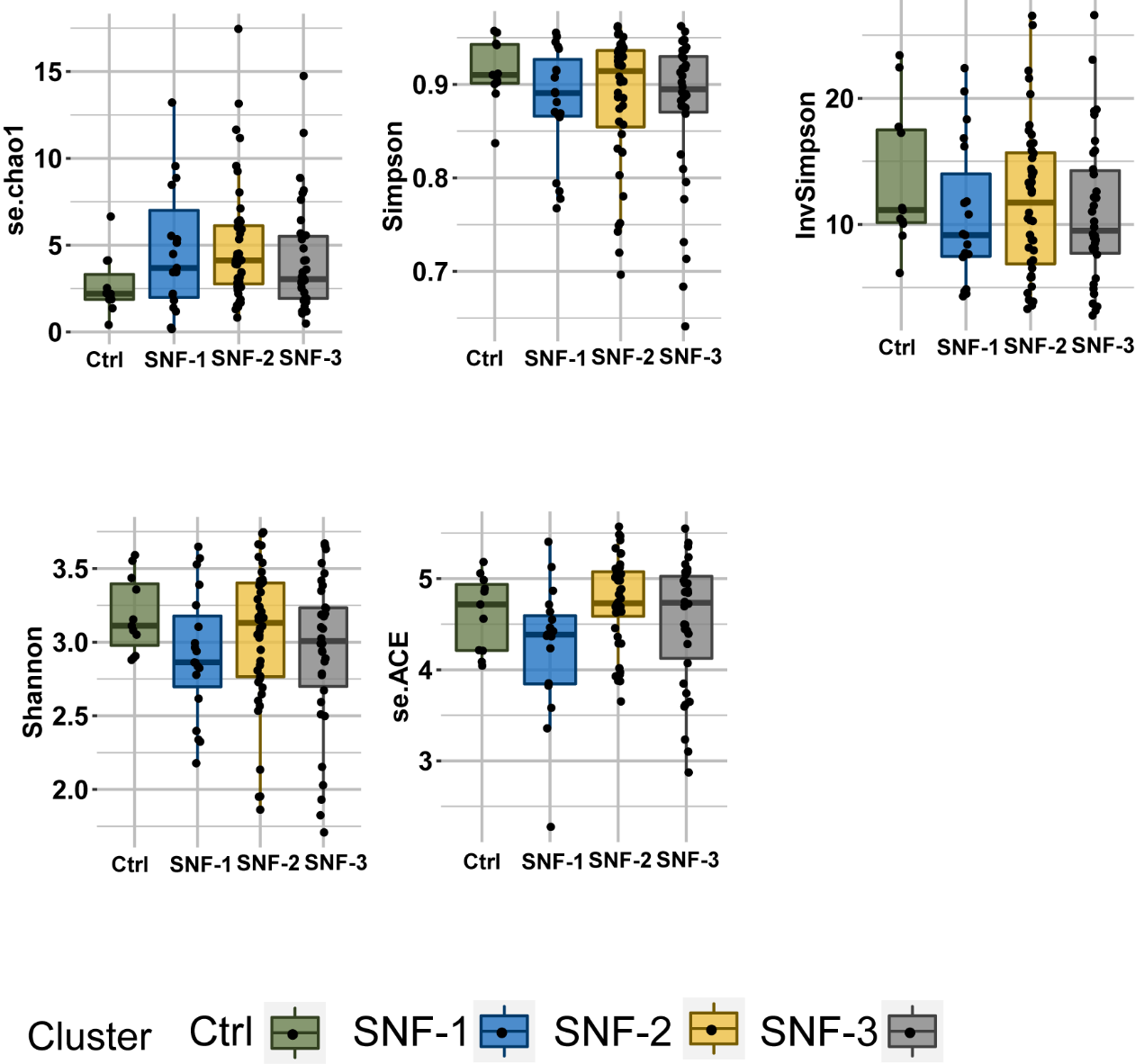

**Fig S3.** Boxplots of alpha diversity indices (se.chao1,Simpson, Shannon, se.ACE, InvSimpson) separated by HIV-cluster. Color is based on groups (Ctrl = green, SNF-1 = blue, SNF-2 = yellow, SNF-3 = grey).

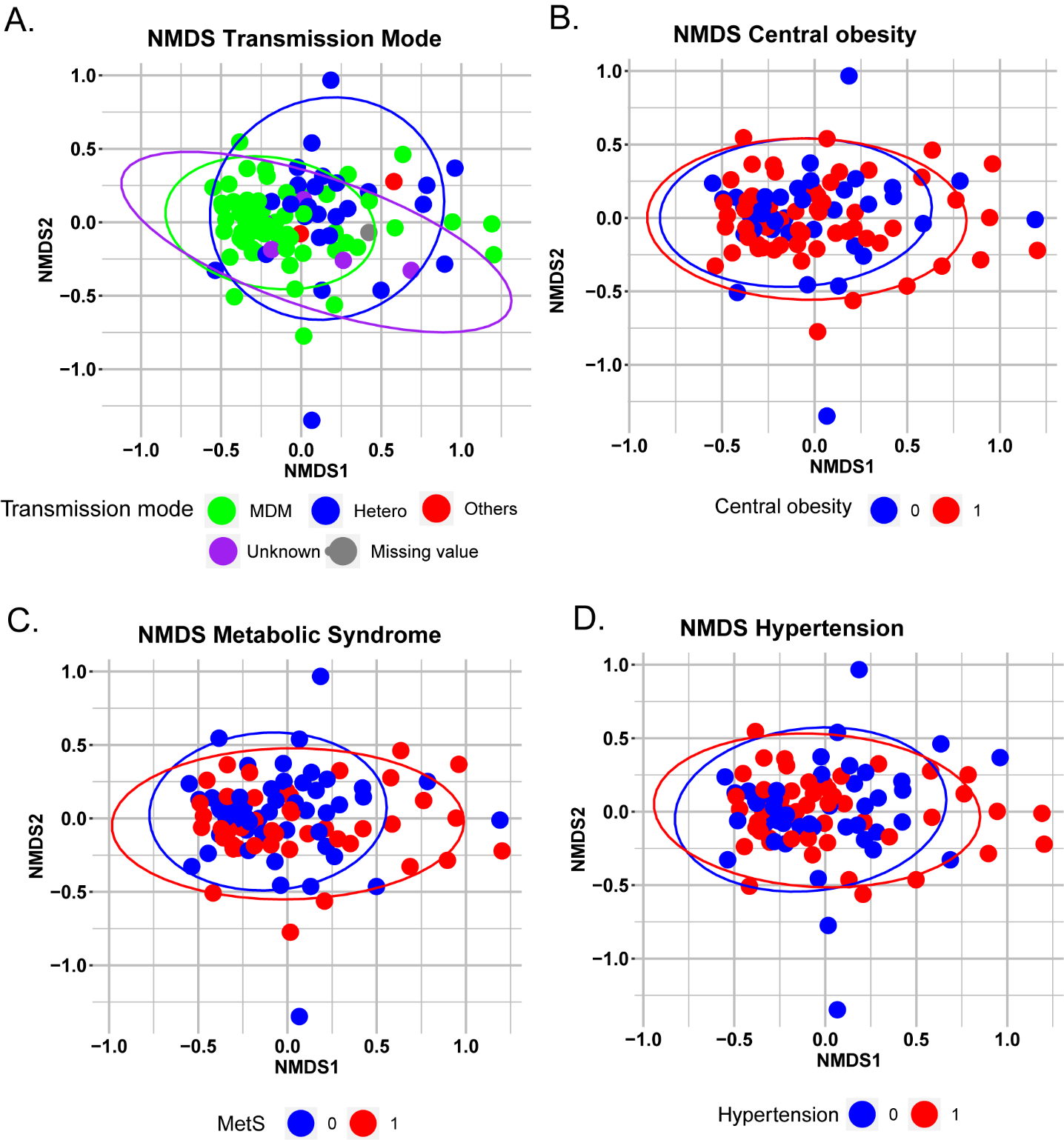
**Fig S4.** Non-metric multidimensional scaling (NMDS) plot of Bray-Curtis distances. Samples are colored by A) Transmission mode B) Central obesity C) Metabolic Syndrome D) Hypertension.

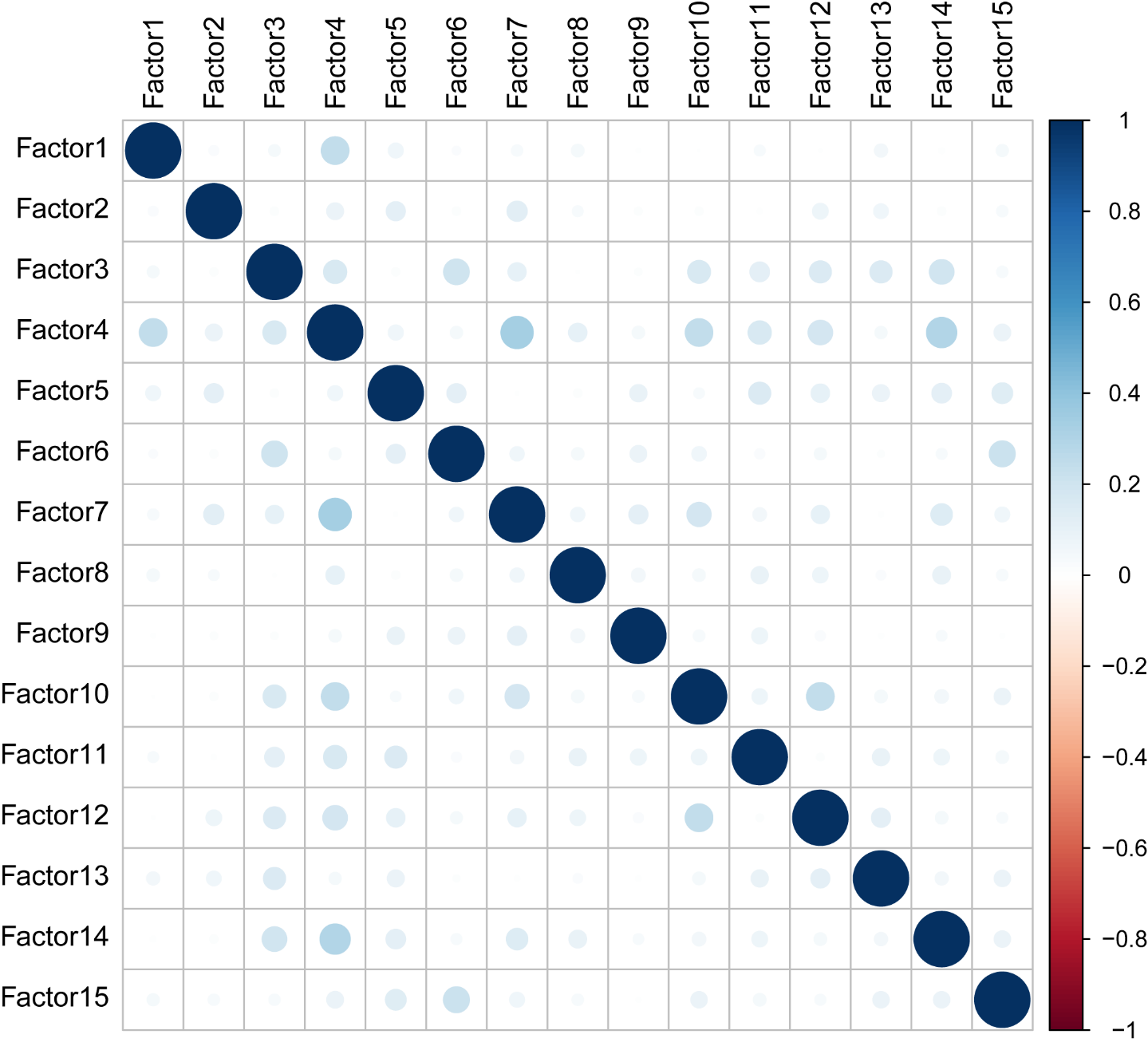

**Fig S5.** Correlation matrix of MOFA factors. Size and transparency are proportional to absolute coefficient of correlation. Color is displayed as a gradient-based coefficient of correlation from -1 (red) to 1 (blue).

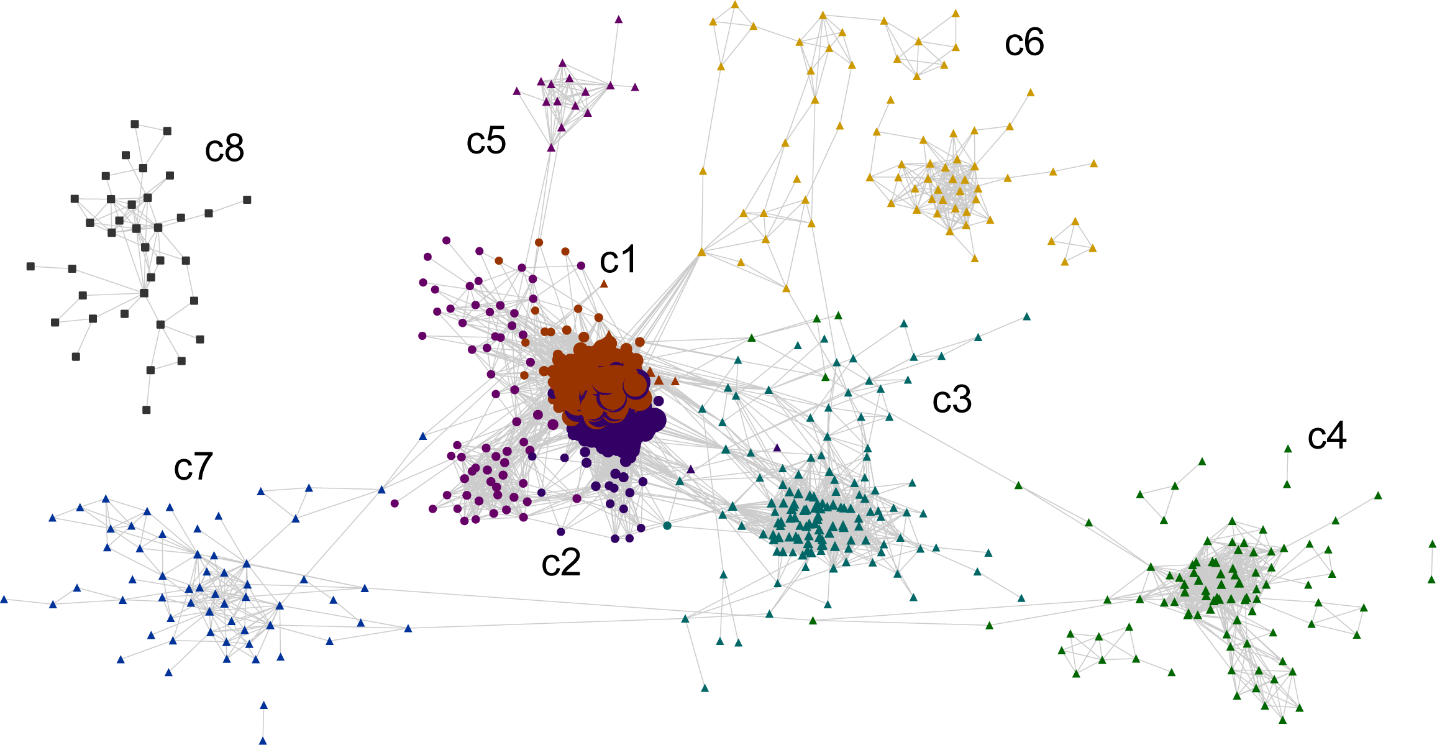

**Fig S6.** Cytoscape consensus co-expression network. Color and label are based on communities.
